# Validation and invalidation of SARS-CoV-2 papain-like protease inhibitors

**DOI:** 10.1101/2021.11.04.467342

**Authors:** Chunlong Ma, Jun Wang

## Abstract

SARS-CoV-2 encodes two viral cysteine proteases, the main protease (M^pro^) and the papain-like protease (PL^pro^), both of which are validated antiviral drug targets. The PL^pro^ is involved in the cleavage of viral polyproteins as well as immune modulation through removing ubiquitin and interferon-stimulated gene product 15 (ISG15) from host proteins. Therefore, targeting PL^pro^ might be a two-pronged approach. Several compounds including YM155, cryptotanshinone, tanshinone I, dihydrotanshinone I, tanshinone IIA, SJB2-043, 6-thioguanine, and 6-mercaptopurine were recently identified as SARS-CoV-2 PL^pro^ inhibitors through high-throughput screening. In this study, we aim to validate/invalidate the reported PL^pro^ inhibitors using a combination of PL^pro^ target specific assays including enzymatic FRET assay, thermal shift binding assay (TSA), and the cell based FlipGFP assay. Collectively, our results showed that all compounds tested either did not show binding or led to denaturation of the PL^pro^ in the TSA binding assay, which might explain their weak enzymatic inhibition in the FRET assay. In addition, none of the compounds showed cellular inhibition of PL^pro^ as revealed by the FlipGFP assay. Therefore, more efforts are needed to search for specific and potent SARS-CoV-2 PL^pro^ inhibitors.

## 1. Introduction

The COVID-19 pandemic is a timely reminder for the urgent need of antivirals, especially broad-spectrum antivirals that could be used as the first line defense against not only current pandemic but also future pandemics.^1^ SARS-CoV-2, together with SARS-CoV and MERS-CoV are the three coronaviruses in the β-coronavirus family that caused pandemic/epidemic in human.^2^ The SARS-CoV the MERS-CoV had higher mortality rates than SARS-CoV-2.^3^ However, SARS-CoV-2 has a much higher transmission rate than SARS-CoV and MERS-CoV, which leads to far greater infection cases and death tolls.

SARS-CoV-2 shares 86% sequence identity with SARS-CoV, which renders the rapid understanding of its viral pathogenicity feasible. SARS-CoV-2 expresses two viral proteases during the viral replication, the main protease (M^pro^; nsp5) and the papain-like protease (PL^pro^; nsp3). Both are cysteine proteases and have been validated as antiviral drug targets.^4-6^ PL^pro^ and M^pro^ cleave the viral polyproteins pp1a and pp1ab at 3 and more than 11 sites, respectively, resulting in individual functional viral proteins for the assembly of viral replication complex. Compared to PL^pro^, M^pro^ is a more amenable drug target and has been the central focus of COVID-19 antiviral discovery. Structurally disparate compounds have been reported as M^pro^ inhibitors from either drug repurposing screening or rational design.^7-8^ Although a large number of reported M^pro^ inhibitors were later proven to be promiscuous cysteine modifiers,^9-13^ several M^pro^ inhibitors have been validated as specific inhibitors and have shown *in vivo* antiviral efficacy in animal model studies.^14-18^ Significantly, two Pfizer M^pro^ inhibitors PF-07304814 and PF-07321332 are advanced to human clinical trials.^17-18^ PL^pro^ is second viral cysteine protease that cleaves the viral polypeptide at three different sites during viral replication. In addition, PL^pro^ modulates host immune response by cleaving ubiquitin and ISG15 (interferon-induced gene 15) from host proteins.^19-21^ In contrast to M^pro^ inhibitors, the development of PL^pro^ inhibitors is still at its infancy.^22^ The most promising PL^pro^ inhibitors are the naphthalene-based GRL0617 series of compounds (Figure 1A).^6, 21, 23-25^ However, their antiviral potency and pharmacokinetic properties need to be further optimized for the *in vivo* animal model study. PL^pro^ specifically recognize the P1-P4 sequence Gly-Gly-X-Leu that is conserved among the nsp1/2, nsp2/3, and nsp3/4 cleavage sites at the viral polyprotein.^26^ The featureless P1 and P2 binding pockets present a grand challenge in developing potent PL^pro^ inhibitors.^26^ As the drug-binding site is far away from the catalytic cysteine (> 10Å), majority of the reported PL^pro^ inhibitors are non-covalent inhibitors.^7^ Although peptidomimetic covalent PL^pro^ inhibitors have been reported, no antiviral activity was shown.^26^ To identify additional novel PL^pro^ inhibitors, several high-throughput screenings have been performed and structurally disparate compounds were found to inhibit PL^pro^ (Figure 1B).^27-30^ To evaluate whether any of the identified candidates might worth further pursuing, we are interested in validating/invalidating the reported PL^pro^ inhibitors using a combination of orthogonal assays including the activity-based FRET enzymatic assay, the thermal shift binding (TSA) assay, and the cell-based FlipGFP assay. Collectively, in contrast to the literature reported results, our data showed that the examined non-GRL0617-based PL^pro^ inhibitors (Figure 1B) had weak enzymatic inhibition and no cellular PL^pro^ inhibition, therefore they should not be referenced as PL^pro^ inhibitors.

**Figure 1.**
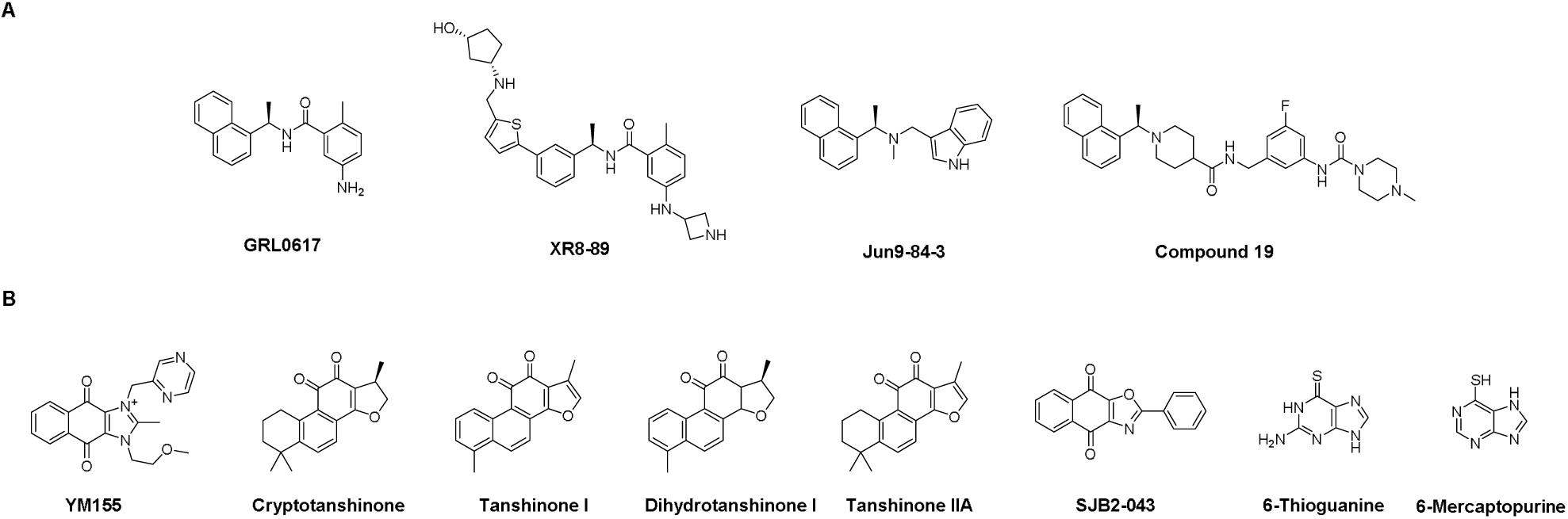
SARS-CoV-2 PL^pro^ inhibitors. (A) GRL0617 analogs. (B) Structurally disparate PL^pro^ inhibitors identified from high-throughput screening or drug repurposing.

## 2. Results

Through a high-throughput screening of over 6,000 bioactive compounds using a quenched fluorescent substrate RLRGG-AMC, Zhao *et al* identified four compounds YM155, cryptotanshinone, tanshinone I, and GRL0617 as potent PL^pro^ inhibitors with IC_50_ values from 1.39 to 5.63 µM.^29^ In plaque reduction assay, YM155, cryptotanshinone, and tanshinone l inhibited SARS-CoV-2 replication in Vero E6 cells with EC_50_ values of 0.17, 0.70, and 2.26 µM, respectively. The X-ray crystal structure of SARS-CoV-2 PL^pro^ with YM155 was solved (PDB: 7D7L), revealing three different binding sites in the thumb domain, the zinc-finger motif, and the substrate-binding pocket. In the substrate binding pocket, YM155 induces a conformational change of Y268 on the BL2 loop and forms a π-stacking interaction, which is similar to the binding mode of GRL0617. Binding at the thumb domain likely impedes the interaction between PL^pro^ and ISG15. The third binding site at the zinc-finger domain might perturb its stability, however, this mechanism remains to be validated.

In our validation study, the positive control GRL0617 showed similar enzymatic inhibition with an IC_50_ of 1.67 µM (Figure 2A). However, we found that the enzymatic inhibition potency for YM155, cryptotanshinone, and tanshinone I against PL_pro_ were ∼10-fold less active than previously reported and the IC_50_ values from our study were 20.16, 52.24, and 18.58 µM, respectively (Figure 2A and Table 2). The discrepancy might be caused by the different substrates used. In Zhao *et al*’s study, RLRGG-AMC was used, while we used a FRET substrate Dabcyl-FTLRGG/APTKV-Edans which spans both the P and P’ sites. Nevertheless, in the thermal shift binding assay, YM155 had no effect on the stability of PL^pro^, while cryptotanshinone and tanshinone I caused destabilization of the protein (Figure 2B and Table 2). In the cell-based FlipGFP assay,^6^ the result for YM155 was not conclusive as it was cytotoxic to the 293T cells (Figure 2C). Both cryptotanshinone and tanshinone I were not active (EC_50_ > 60 µM) (Figure 2C). Overall, our results indicate that cryptotanshinone and tanshinone I were not specific PL^pro^ inhibitors, while YM155 might be a specific PL^pro^ inhibitor. However, the large discrepancy between the enzymatic inhibition (IC_50_ = 20.16 µM) and the cellular antiviral activity of YM155 (EC_50_ = 0.17 µM) suggest that the other mechanisms might contribute to its potent antiviral activity against SARS-CoV-2.

**Figure 2.**
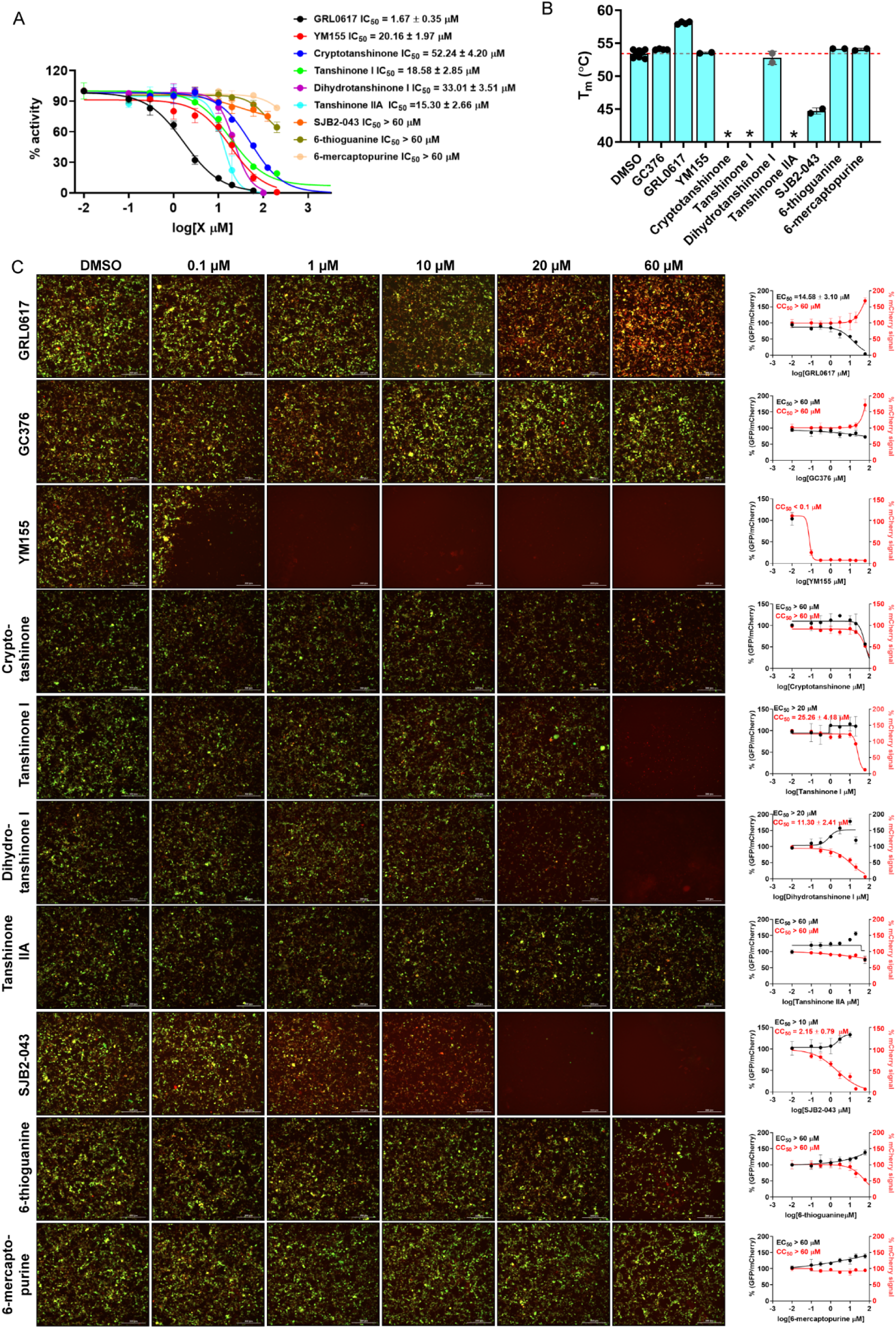
Pharmacological characterization of SARS-CoV-2 PL^pro^ inhibitors. (A) Enzymatic inhibitory activity against SARS-CoV-2 PL^pro^ in FRET-based assay. (B) Thermal shift assay of the SARS-CoV-2 PL^pro^ inhibitors in stabilizing the SARS-CoV-2 PL^pro^. * means melting peak was not observed the in the presence of the inhibitor. The dashed red line indicates the mean of SARS-CoV-2 PL^pro^ T_m_ in the absence of testing inhibitors. (C) Cell-based FlipGFP assay for the quantification of the cellular activity of SARS-CoV-2 PL^pro^ inhibitors. Representative images of FlipGFP PL^pro^ assay with the positive control GRL0617 and the negative control GC-376. Dose™response curves of the ratio of GFP/mCherry fluorescence with SARS-CoV-2 PL^pro^ inhibitors were plotted at right side column; mCherry signal alone was used to normalize protein expression level or calculate compound cytotoxicity.

In another study, cryptotanshinone, together with two analogs dihydrotanshinone I and tanshinone IIA were identified as SARS-CoV-2 PL^pro^ inhibitors through a high-throughput screening (HTS).^28^ Cryptotanshinone, dihydrotanshinone I and tanshinone IIA inhibited SARS-CoV-2 PL^pro^ with IC_50_ values of 1.336, 0.5861, and 1.571 µM, respectively. It is noted that no complete inhibition was observed for cryptotanshinone, therefore the IC_50_ value might not be trustworthy. To rule out the possibility of fluorescence interference, these compounds were also tested in the gel-based PL^pro^ assay using the GST-nsp2/3-MBP substrate. All these compounds inhibited the digestion of the protein substrate with tanshinone IIA and cryptotanshinone showing higher potency than dihydrotanshinone I. Dihydrotanshinone I also inhibited the deubiquitinase and deISGlase activities of PL^pro^ in the gel-based digestion assay. In the SARS-CoV-2 antiviral assay in Vero E6 cells, dihydrotanshinone I had the highest potency with an EC_50_ of 8.148 µM, while cryptotanshinone and tanshinone I were not active (EC_50_ > 200 µM). The lack of correlation between the enzymatic inhibition and the cellular antiviral activity might due to cell membrane permeability or off-target effects.

In our validation study, both dihydrotanshinone I and tanshinone IIA were greater than 10-fold less potent in the enzymatic assay with IC_50_ values of 33.01 and 15.30 µM, respectively (Figure 2A and Table 2). Dihydrotanshinone I did not bind to PL^pro^ as shown by the results from the TSA assay (Figure 2B). Tanshinone IIA led to denaturation of the protein (no melting peak). Dihydrotanshinone I and tanshinone IIA were not active in the FlipGFP assay (IC_50_ > 20 µM) (Figure 2C). Overall, it appears that dihydrotanshinone I and tanshinone IIA were not specific SARS-CoV-2 PL^pro^ inhibitors, and the antiviral activity of dihydrotanshinone I might involve other mechanisms.

Cho *et al* reported SJB2-043 a SARS-CoV-2 PL^pro^ inhibitor through a focused screening of a library of deubiquitinase inhibitors.^27^ SJB2-043 did not achieve complete inhibition and had an apparent IC_50_ of 0.56 µM. When Ub-AMC was used as a substrate, the IC_50_ of SJB2-043 was 0.091 µM. Similarly, no complete inhibition was achieved. In comparison, the positive control GRL0617 showed complete inhibition when both Z-LRGG-AMC and Ub-AMC were used as substrates, and the IC_50_ values were 1.37 and 1.80 µM, respectively. Molecular docking suggests that SJB2-043 binds to an allosteric site in PL^pro^.

When repeated in our assay, SJB2-043 did not inhibit the SARS-CoV-2 PL^pro^ (IC_50_ > 60 µM) (Figure 2A). The discrepancy might be caused by different methods used for data fitting. In our assay, SJB2-043 did not achieve more than 50% inhibition at the highest concentration tested, which is 60 µM, therefore we deemed the IC_50_ greater than 60 µM. In the TSA binding assay, SJB2-043 led to the destabilization of PL^pro^ (ΔT_m_ = -8.72 µM) (Figure 2B). The result from the FlipGFP assay was not conclusive as SJB2-043 was cytotoxic (Figure 2C). Overall, SJB2-043 might not be a specific SARS-CoV-2 PL^pro^ inhibitor and only shows partial inhibition.

Swaim *et al* reported that 6-thioguanine inhibited SARS-CoV-2 replication in Vero E6 cells with an EC_50_ of 2.13 µM.^30^ Mechanistic studies showed that 6-thioguanine had potent inhibition against PL^pro^ in cells and *in vitro* using the TAP-nsp123, TAP-nsp23, and Pro-ISG15-HA substrates. Another study from Fu *et al* showed that 6-thioguanine inhibited SARS-CoV-2 PL^pro^ in the enzymatic assay with an IC_50_ of 72 µM.^31^ In our study, 6-thioguanine was not active in the FRET-based enzymatic assay (IC_50_ > 60 µM) (Figure 2A). 6-thioguanine also did not show binding in the TSA assay nor inhibition in the FlipGFP assay (Figures 2B and 2C). As such, the antiviral activity of 6-thioguanine might involve other mechanisms. 6 mercaptopurine, which is an analog of 6-thioguanine, was similar not active in all three assays (Figure 2 and Table 2). Therefore, it can be concluded that 6-thioguanine and 6-mercaptopurine were not SARS-CoV-2 PL^pro^ inhibitors.

## 3. Discussion

Drug repurposing is often viewed as a fast-track drug discovery approach since it can potentially bypass the length safety tests before entering clinical trials.^32^ However, we should remain cautiously optimistic about this approach.^33^ Since the existing bioactive compounds were not specifically designed and optimized for the screening drug target, the identified hits need to be vigorously characterized to validate the target specificity. The SARS-CoV-2 M^pro^ and PL^pro^ are cysteine proteases which are prone to non-specific inhibition by alkylating agents or re-dox cycling compounds. Indeed, our recent studies, together with others, have collectively shown that a number of reported M^pro^ inhibitors including ebselen, carmofur, disulfiram, and shikonin are promiscuous non-specific cysteine protease inhibitors.^9-13^ With our continuous interest in validation/invalidation of literature reported M^pro^ and PL^pro^ inhibitors, in this study we characterized eight PL^pro^ inhibitors using a combination of FRET-based enzymatic assay, TSA binding assay, and the cell-based FlipGFP PL^pro^ assay. It is expected that specific inhibitors will show consistent results from all three assays. Among the compounds tested, cryptotanshinone, tanshinone I, dihydrotanshinone I, and tanshinone IIA were also previously reported as potent SARS-CoV PL^pro^ inhibitors with IC_50_ values ranging from 0.8 to 8.8 µM.^34^ Interesting, they also inhibited SARS-CoV M^pro^ with IC_50_ values ranging from 14.4 to 226.7 µM. Since M^pro^ and PL^pro^ do not share structural and sequence similarities, the dual inhibitory activities of these tanshinones, coupled with the weak PL^pro^ enzymatic inhibition from our study, suggest that they might have a promiscuous mechanism of action.

Similarly, 6-thioguanine and 6-mercaptopurine were previously reported as SARS-CoV PL^pro^ inhibitors with IC_50_ values of 5.0 and 21.6 µM, respectively.^35-36^ In addition, Cheng *et al* reported that 6-thioguanine and 6-mercaptopurine are competitive inhibitors of MERS-CoV PL^pro^ with IC_50_ values of 24.4 and 26.9 µM, respectively.^37^ In contrast, YM155 and SJB2-043 were not previously shown to inhibit SARS-CoV PL^pro^. Given the 82.6% sequence similarity between SARS-CoV-2 and SARS-CoV PL^pro^s, it appears not a surprising finding that cryptotanshinone, tanshinone I, dihydrotanshinone I, tanshinone IIA, and 6-thioguanine were also reported as SARS-CoV-2 PL^pro^ inhibitors from recent studies. Nevertheless, our independent study has shown that all eight compounds had drastically reduced enzymatic inhibition against PL^pro^ compared to reported values (Table 1). In addition, all compounds either had no effect or destabilize PL^pro^ as shown by the TSA binding assay (Table 1). Furthermore, cryptotanshinone, tanshinone I, dihydrotanshinone I, tanshinone IIA, 6-thioguanine, and 6-mercaptopurine were not active in the FlipGFP PL^pro^ assay, suggesting there is a lack of cellular PL^pro^ target engagement. YM155 and SJB2-043 were cytotoxic, therefore it remains unknown whether they can selectively bind to PL^pro^ inside the cell. Overall, our study calls for more stringent validation of the reported SARS-CoV-2 PL^pro^ inhibitors to avoid the failure in the follow up lead optimization and translational development.

**Table 1.**
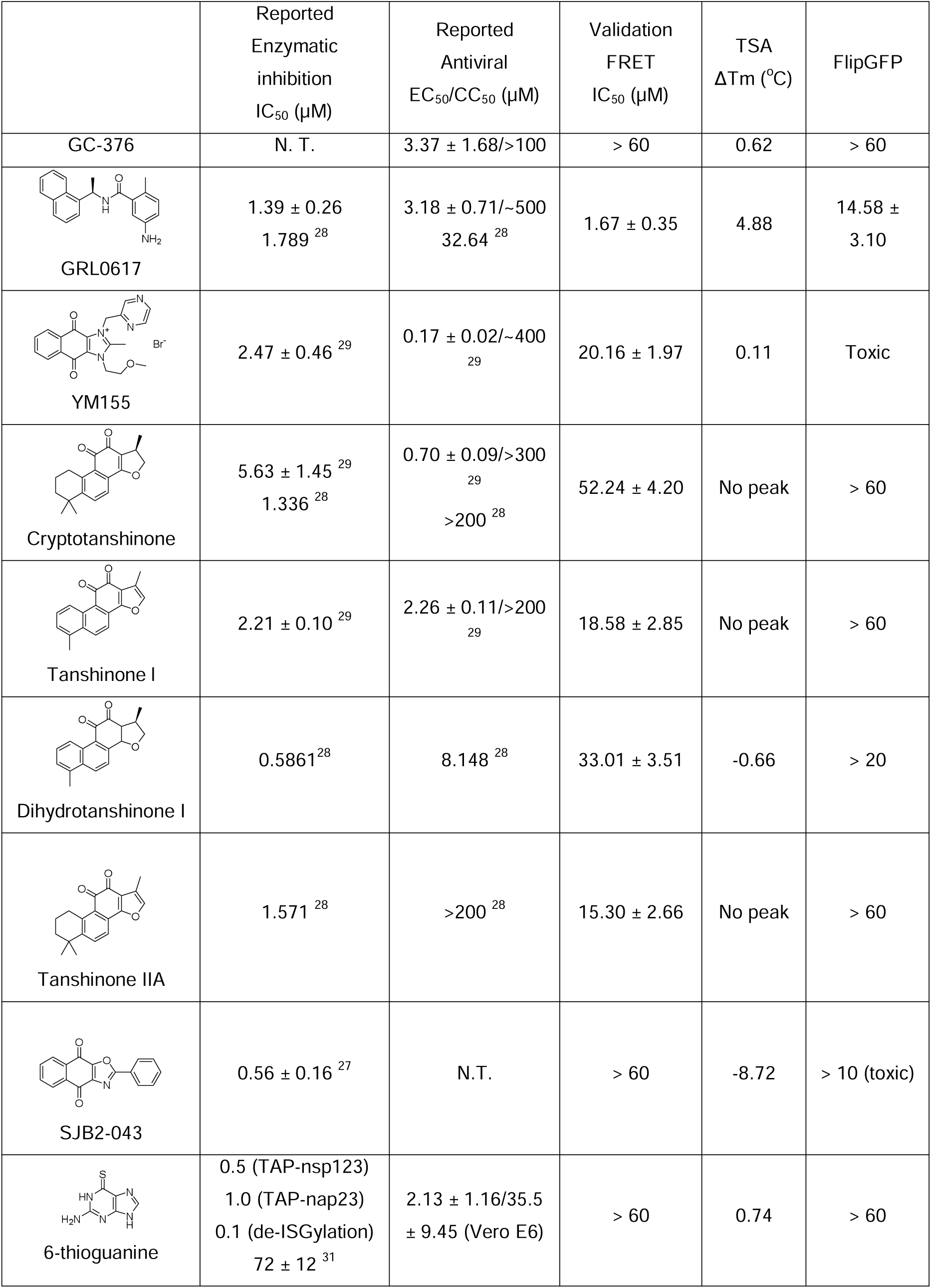
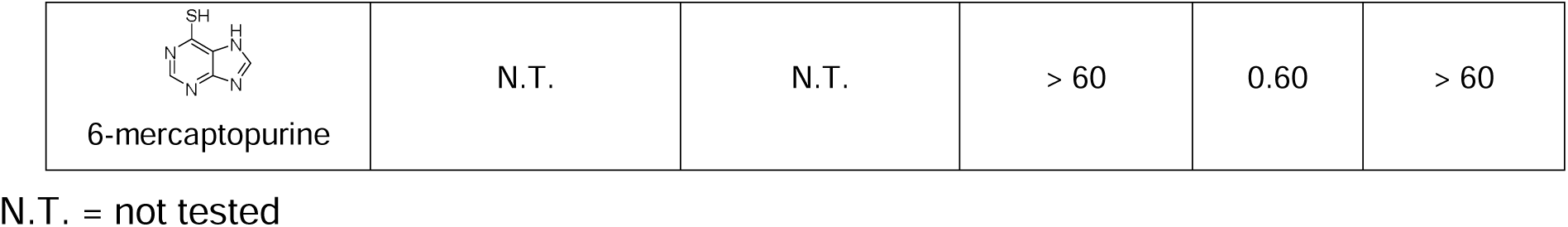
Validation/invalidation of SARS-CoV-2 PL^pro^ inhibitors.

|  | Reported<br>Enzymatic<br>inhibition<br>IC<sub>50</sub> (μM) | Reported<br>Antiviral<br>EC<sub>50</sub>/CC<sub>50</sub> (μM) | Validation<br>FRET<br>IC<sub>50</sub> (μM) | TSA<br>ΔT<sub>m</sub> (°C) | FlipGFP |
| --- | --- | --- | --- | --- | --- |
| GC-376 | N. T. | 3.37 ± 1.68/>100 | > 60 | 0.62 | > 60 |
| GRL0617 | 1.39 ± 0.26<br>1.789<sup>28</sup> | 3.18 ± 0.71/~500<br>32.64<sup>28</sup> | 1.67 ± 0.35 | 4.88 | 14.58 ±<br>3.10 |
| YM155 | 2.47 ± 0.46<sup>29</sup> | 0.17 ± 0.02/~400<sup>29</sup> | 20.16 ± 1.97 | 0.11 | Toxic |
| Cryptotanshinone | 5.63 ± 1.45<sup>29</sup><br>1.336<sup>28</sup> | 0.70 ± 0.09/>300<sup>29</sup><br>>200<sup>28</sup> | 52.24 ± 4.20 | No peak | > 60 |
| Tanshinone I | 2.21 ± 0.10<sup>29</sup> | 2.26 ± 0.11/>200<sup>29</sup> | 18.58 ± 2.85 | No peak | > 60 |
| Dihydrotanshinone I | 0.5861<sup>28</sup> | 8.148<sup>28</sup> | 33.01 ± 3.51 | -0.66 | > 20 |
| Tanshinone IIA | 1.571<sup>28</sup> | >200<sup>28</sup> | 15.30 ± 2.66 | No peak | > 60 |
| SJB2-043 | 0.56 ± 0.16<sup>27</sup> | N.T. | > 60 | -8.72 | > 10 (toxic) |
| 6-thioguanine | 0.5 (TAP-nsp123)<br>1.0 (TAP-nap23)<br>0.1 (de-ISGylation)<br>72 ± 12<sup>31</sup> | 2.13 ± 1.16/35.5<br>± 9.45 (Vero E6) | > 60 | 0.74 | > 60 |
| 6-mercaptapurine | N.T. | N.T. | > 60 | 0.60 | > 60 |

N.T. = not tested

## 4. Materials and methods

### Chemicals and peptide substrate

The SARS-CoV-2 PL^pro^ substrate Dabcyl-FTLRGG/APTKV-Edans was synthesized by the solid-phase synthesis and the detailed procedure was described in our previous publication.^6, 10^ The testing compounds were ordered from the following sources: tanshinone I (Sigma T5330), dihydrotanshinone I (Sigma D0947), tanshinone IIA (Sigma SML2517), cryptotanshinone (Sigma C5624), YM-155 (APExBIO A4221), and SJB2-043 (APExBIO A3823).

### SARS-CoV-2 PL^pro^ expression and purification

SARS-CoV-2 papain-like protease (PL^pro^) gene (ORF 1ab 1564–1876) from strain BetaCoV/Wuhan/WIV04/2019 with *E. coli* codon optimization was ordered from GenScript in the pET28b(+) vector. The detailed expression and purification procedures were previously described.^6, 10^

### FRET-Based enzymatic assay

For the IC_50_ measurement with FRET-based assay, the reaction was carried out in 96-well format with100 µl of 200 nM PL^pro^ protein in a PL^pro^ reaction buffer (50 mM HEPES pH 7.5, 5 mM DTT and 0.01% Triton X-100); 1 µl of testing compounds at various concentration was added to each well and was incubated at 30 °C for 30 min. The reaction was initiated by adding 1 µl of 1 mM FRET substrate and was monitored in a Cytation 5 image reader with filters for excitation at 360/40 nm and emission at 460/40 nm at 30 °C for 1 h. The initial velocity of the enzymatic reaction was calculated from the initial 10 min enzymatic reaction. The IC_50_ was calculated by plotting the initial velocity against various concentrations of testing compounds by use of a four parameters variable slope dose-response curve in Prism 8 software.

### Differential Scanning Fluorimetry (DSF)

The thermal shift binding assay (TSA) was carried out using a Thermo Fisher QuantStudio 5 Real-Time PCR system as described previously.^6, 10^

### Cell-Based FlipGFP-PL^pro^ Assay

Cell-based FlipGFP-PL^pro^ assay was established as previously described. Briefly, 293T cells were maintained in Dulbecco’s modified Eagle’s medium (DMEM), supplemented with 10% heat-inactivated FBS and 1% penicillin-streptomycin antibiotics in a 37 °C incubator with 5% CO_2_. 96-well Greiner plate (catalog no. 655090) was seeded with 293T cells to overnight 70–90% confluency. 50 ng of pcDNA3-flipGFP-PLpro-T2A-mCherry plasmid and 50 ng of pcDNA3.1-SARS2-PL^pro^ were used each well in the presence of transfection reagent TransIT-293 (Mirus) per manufacturer protocol. Three hours after transfection, 1 μl of testing compound was added to each well at 100-fold dilution. 2 days after transfection images were obatined with a Cytation 5 imaging reader (Biotek) with GFP and mCherry channels. SARS-CoV-2 PL^pro^ protease activity was evaluated by the ratio of GFP signal intensity over the mCherry signal intensity. The compound FlipGFP-PLP assay IC_50_ value was calculated by plotting the GFP/mCherry signal to the testing compound concentration with a four-parameter variable slope dose–response function in Prism 8. The mCherry signal alone was utilized to determine the compound cytotoxicity.

## Author Contributions

C. M. performed the enzymatic assay, thermal shift binding assay, and the FlipGFP assay. J. W. designed and supervised this study. J. W. wrote the manuscript with contribution from C. M.

## Acknowledgements

This research was supported by the National Institutes of Health (NIH) (Grant AI147325, AI157046, and AI158775) and the Arizona Biomedical Research Centre Young Investigator grant (ADHS18-198859) to J. W.

## For Table of Contents Use Only

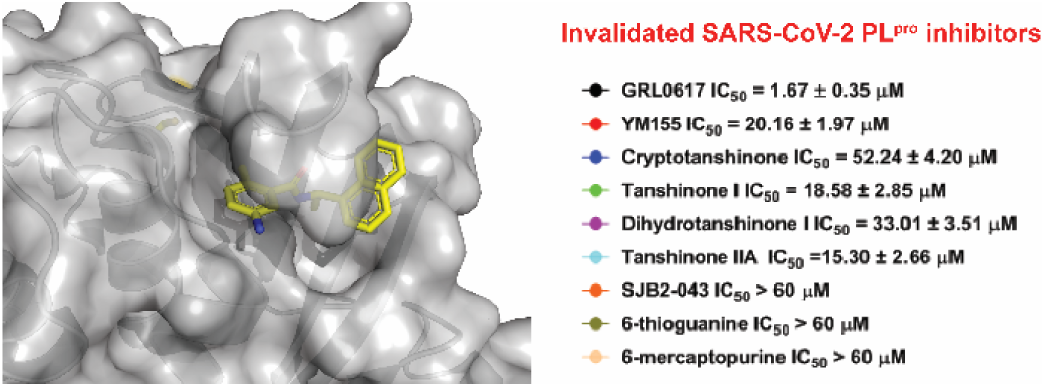

